# Acid-Sensing Ion Channel 1a Deficiency Drives Endocrine Hypertension in Male Mice

**DOI:** 10.1101/2025.03.25.645371

**Authors:** Selina M. Garcia, Megan N. Tuineau, Xavier A. DeLeon, Neil D. Detweiler, Shristey Tamang, Nancy L. Kanagy, Laura V. Gonzalez Bosc, Thomas C. Resta, Jay S. Naik, Nikki L. Jernigan

## Abstract

**Background:** Acid-sensing ion channel 1a (ASIC1a) is an H^+^-gated cation channel that responds to extracellular acidosis in both normal and pathological states, including ischemia, inflammation, and metabolic disturbances. While ASIC1a regulates vascular reactivity, its role in blood pressure regulation remains unclear, particularly concerning sex, aging, and disease. This study aims to investigate whether ASIC1a: 1) contributes to cardiovascular function in a sex-dependent manner; 2) plays a dynamic role in cardiovascular homeostasis with aging; and 3) modulates the development of angiotensin II-induced systemic hypertension.

**Methods:** Radiotelemeters were implanted in 6- and 18-month-old male and female wild-type (*Asic1a*^+/+^) and ASIC1a knockout (*Asic1a*^-/-^) mice to monitor mean arterial blood pressure and heart rate under baseline conditions and in response to angiotensin II. Blood gases, electrolytes, hormones, and end-organ injury were also assessed.

**Results:** Aged male *Asic1a*^-/-^ mice develop hypertension driven by aldosterone excess and sympathetic overactivity, which is accompanied by cardiac hypertrophy, aortic fibrosis, and glomerular hypertrophy. Female *Asic1a*^-/-^ mice remain unaffected. In male *Asic1a*^-/-^ mice, hyperaldosteronism occurs independent of the renin-angiotensin system and mitigates angiotensin II-induced hypertension. Furthermore, 6-month-old male *Asic1a*^-/-^ mice exhibit elevated corticosterone, hypokalemia, reduced urine osmolality, increased pulse pressure, and cardiomyocyte hypertrophy that precedes hypertension.

**Conclusions:** These findings establish ASIC1a as a novel, sex-specific regulator of cardiovascular function, linking early corticosterone excess in male mice to hyperaldosteronism and implicating ASIC1a deficiency as a potential driver of endocrine-related hypertension.

**GRAPHICAL ABSTRACT:** 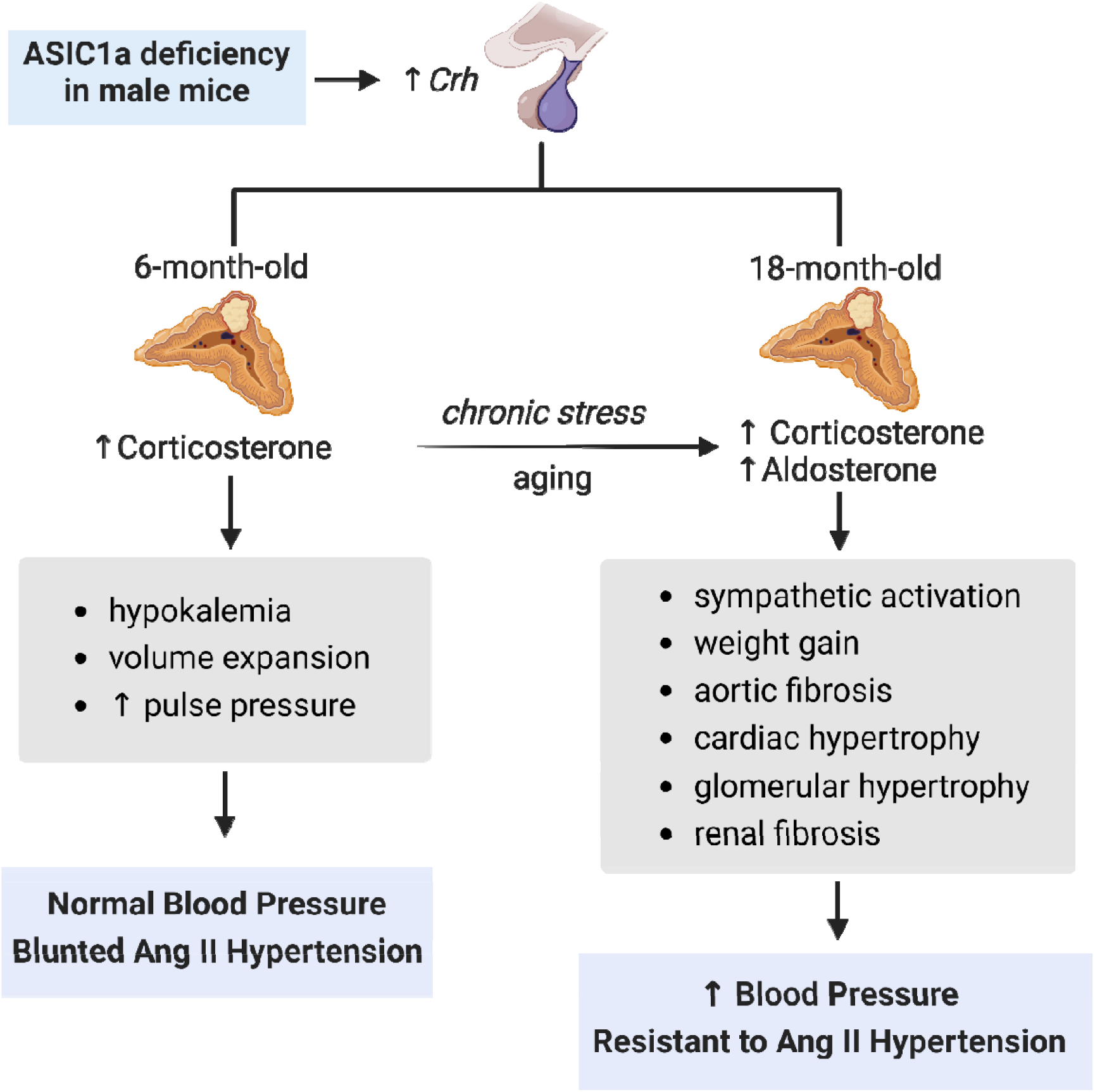

## INTRODUCTION

During pathological conditions, extracellular acidification can occur in response to ischemia, inflammation, and metabolic disturbances—all important contributors to cardiometabolic diseases. While several membrane receptors and ion channels can respond to changes in pH, acid-sensing ion channels (ASIC) are particularly sensitive, with some isoforms having an activation threshold close to pH 7.0^1^. ASICs are voltage-insensitive, H^+^-gated ion channels that are encoded by 5 genes (ACCN1-5), giving rise to 7 ASIC protein subunits/isoforms (1a, 1b, 2a, 2b, 3, 4, and 5)^2^. Three subunits—which can be homomeric or heteromeric—form a functional channel that primarily conducts inward Na^+^ currents, however, homomeric ASIC1a and heteromeric ASIC1a/ASIC2b also conduct Ca^2+ 3–6^. The role of ASIC is best characterized in the central and peripheral nervous systems, where they regulate synaptic plasticity, learning and memory, mechanotransduction, and sensory perception^7,8^. However, ASIC is also expressed in other cell types, including cardiac myocytes, fibroblasts, vascular smooth muscle, and endothelial cells^9–12^.

The role of ASIC in cardiovascular homeostasis was first identified in sensory neurons where ASIC2 functions as the mechanosensor in aortic baroreceptors^13^ and ASIC3 as the acid sensor in carotid body glomus cells^14^. As a result, ASIC2 knockout (*Asic2*^-/-^) mice have elevated mean arterial blood pressure (MABP) and heart rate (HR), whereas ASIC3 knockout (*Asic3*^-/-^) mice exhibit lower MABP compared to wild-type mice^13,15^. While these data provide evidence that ASIC2 and ASIC3 influence cardiovascular function through neural sensory signaling, the role of ASIC1a in cardiovascular homeostasis remains poorly understood, as current findings are largely limited to individual vascular beds.

In the cerebral vasculature, <u>neural</u> ASIC1a mediates vasodilatory responses to hypercapnia^16^. In mesenteric resistance arteries, <u>endothelial</u> ASIC1a facilitates endothelium-dependent dilation by activating small- and intermediate-conductance Ca^2+^-activated K^+^ channel-mediated hyperpolarization^12^. Given its role in vasodilatory responses, one might expect ASIC1a knockout (*Asic1a*^-/-^) mice to exhibit endothelial dysfunction and elevated MABP. However, MABP and HR were reportedly similar in anesthetized wild-type (*Asic1a*^+/+^) and *Asic1a*^-/-^ mice [mixed sex analysis]^17^. This may suggest that ASIC1a regulates blood flow locally within specific vascular beds rather than exerting a broader influence on systemic vascular resistance or blood pressure regulation. Alternatively, ASIC1a may have a minimal effect under normal physiological conditions but play a more significant role during aging or pathological states. Supporting this notion, our previous work indicates that while <u>smooth muscle</u> ASIC1a has a minor impact on pulmonary vascular reactivity under normal conditions, it significantly contributes to hypoxia-induced pulmonary hypertension^11,18^.

To fully elucidate ASIC1a’s role in cardiovascular regulation, we aim to answer three key questions: Does ASIC1a’s contribution to cardiovascular homeostasis change with aging? Does ASIC1a play a role in angiotensin II (Ang II)-induced systemic hypertension? Is ASIC1a’s influence on cardiovascular function sex-dependent? Addressing these questions will provide critical insights into ASIC1a’s function in cardiovascular health and disease.

## METHODS

### Data Availability

The authors declare that all supporting data, analytical methods, and study materials are available within the article and its Data Supplement or can be obtained from the corresponding author upon reasonable request.

### Animal Studies

Studies were completed in male and female, 6- and 18-month-old, wildtype (*Asic1a*^*+/+*^), global ASIC1a knockout (*Asic1a*^*-/-*^), and inducible smooth muscle-specific ASIC1a knockout (MHC^CreER^-*Asic1a* ^fl/fl^) mice. All protocols used in this study abide by the National Institutes of Health guidelines for animals and were reviewed by the Institutional Animal Care and Use Committee of the University of New Mexico School of Medicine (Protocol #22-201292-HSC). Further details regarding the experimental animals are provided in the **Major Resource Table** according to ARRIVE 2.0 guidelines. Detailed experimental methods are provided in the **Data Supplement**.

### Statistical Analysis

Unless otherwise stated, values of n refer to the number of animals in each group. All data are presented as mean ± standard error of the mean (SEM). Statistical comparisons were made using Prism 10 (GraphPad Software), where a probability of <0.05 with a power level of 0.80 was accepted as statistically significant for all comparisons. Additional information for each set of experiments, sample size, and statistical analysis are detailed in the Figure legend.

## RESULTS

### ASIC1a Plays a Paradoxical Role in Blood Pressure Regulation: Protecting Males While Facilitating Age-related Increases in Females

All mice demonstrated a distinct diurnal cycle, with significantly higher MABP and HR during the active dark cycle compared to the resting light cycle (**Figure 1A-B**). In 6-month-old mice, males exhibited higher MABP than females, regardless of genotype (**Figure 1C**). Female *Asic1a*^+/+^ mice showed an age-related increase in MABP and HR, which was absent in female *Asic1a*^-/-^ mice (**Figure 1C-D**). This increase was particularly evident during the resting light cycle, while no significant differences in MABP were observed during the active dark cycle, nor in systolic or diastolic pressures (**Table S1**). In contrast, male *Asic1a*^-/-^ mice experienced a significant age-dependent increase in MABP and HR (**Figure 1C-D**), driven by increases in both systolic and diastolic pressures during both light and dark cycles (**Table S1**). Interestingly, both 6- and 18-month-old male *Asic1a*^-/-^ mice have increased pulse pressure compared to male *Asic1a*^+/+^ mice, a phenomenon not observed in female mice (**Table S1**).

**Figure 1.**
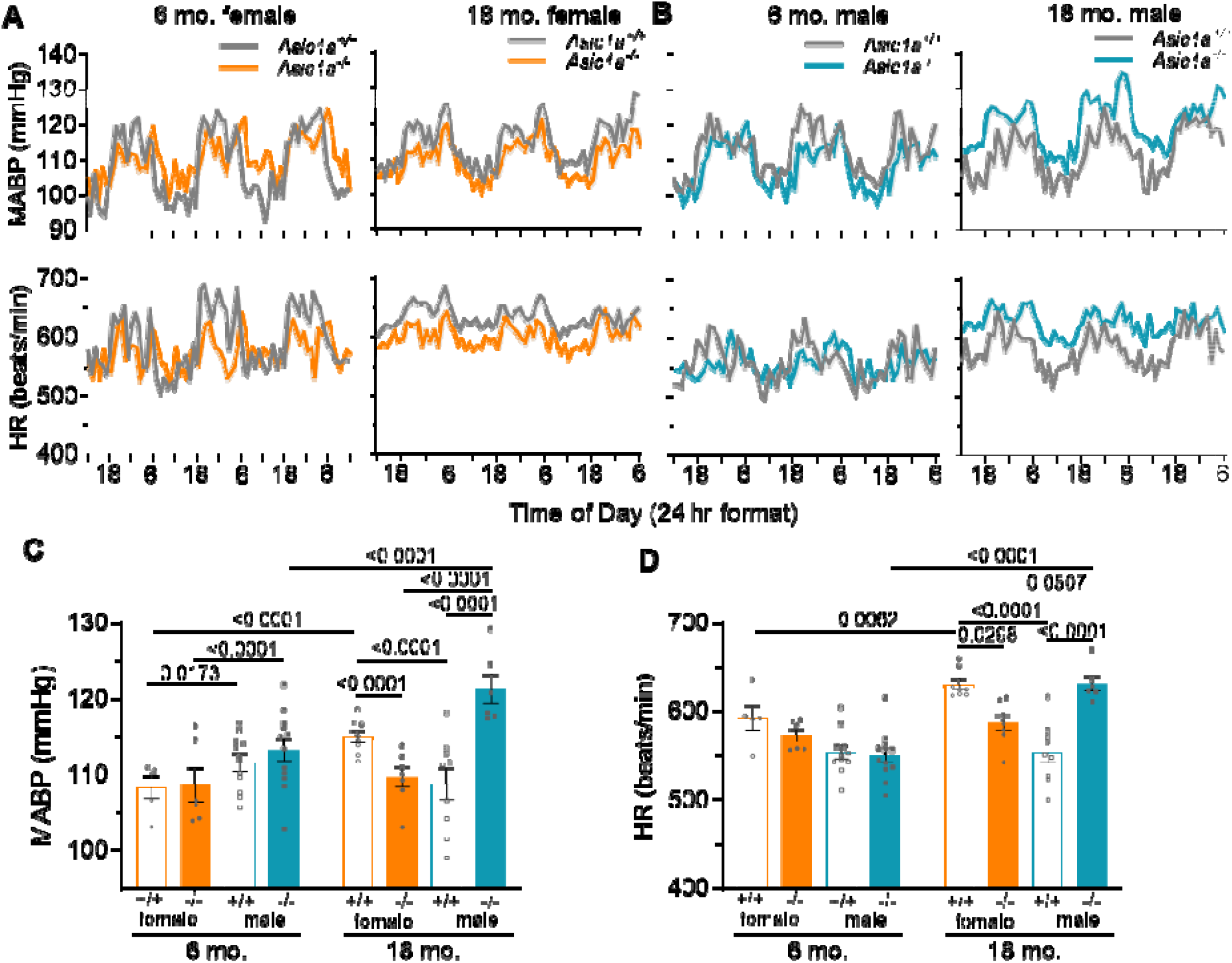
ASIC1a protects male mice but facilitates age-induced increases in blood pressure and heart rate in female mice. Averaged traces and 24-hr summary data showing A, C) mean arterial blood pressure (MABP, mmHg) and B, D) heart rate (HR, beats/min) recorded over 72 hrs in 6- and 18-month-old female (orange) and male (teal) *Asic1a*^+/+^ (open) and *Asic1a*^-/-^ (filled) mice. Light grey boxes indicate the dark (active) cycle. Analyzed by three-way ANOVA, significant interactions between age x sex x genotype [C) p = 0.0011 and D) p = 0.0003] were compared using Tukey’s multiple comparisons tests [* p<0.05 vs age-matched *Asic1a*^+/+^, # p < 0.05 vs corresponding 6-month-old adult, and $ p < 0.05 vs corresponding female]; n = animals/group.

This increase in pulse pressure could be due to increased stroke volume or decreased aortic compliance (i.e. stiffer aorta) in male *Asic1a*^-/-^ mice. Together, these findings demonstrate an age-specific role for ASIC1a in cardiovascular hemodynamics, with a prominent sex-specific difference: ASIC1a deletion protects female mice from age-related increases in MABP and HR, while eliciting hypertension in male mice. Of note, aged *Asic1a*^-/-^ mice weighed significantly more than *Asic1a*^+/+^ mice, regardless of sex, MABP, or HR (**Table S1**).

### Asic1a deletion leads to age-dependent protection against Ang II-induced hypertension in male mice

In addition to defining the effects of ASIC1a deletion on cardiovascular function with age, we examined its effects on Ang II-induced hypertension. In 6-month-old male *Asic1a*^-/-^ mice, increases in MABP in response to Ang II (600 ng/kg/min) were significantly blunted at 21-28 days compared to *Asic1a*^+/+^ mice (**Figure 2A,C**). As *Asic1a*^-/-^ male mice age, they became resistant to Ang II hypertension, showing no significant elevation in MABP relative to baseline measurements (**Figure 2B,C**). Ang II had only modest effects on HR across all groups (**Figure 2D-F**).

**Figure 2.**
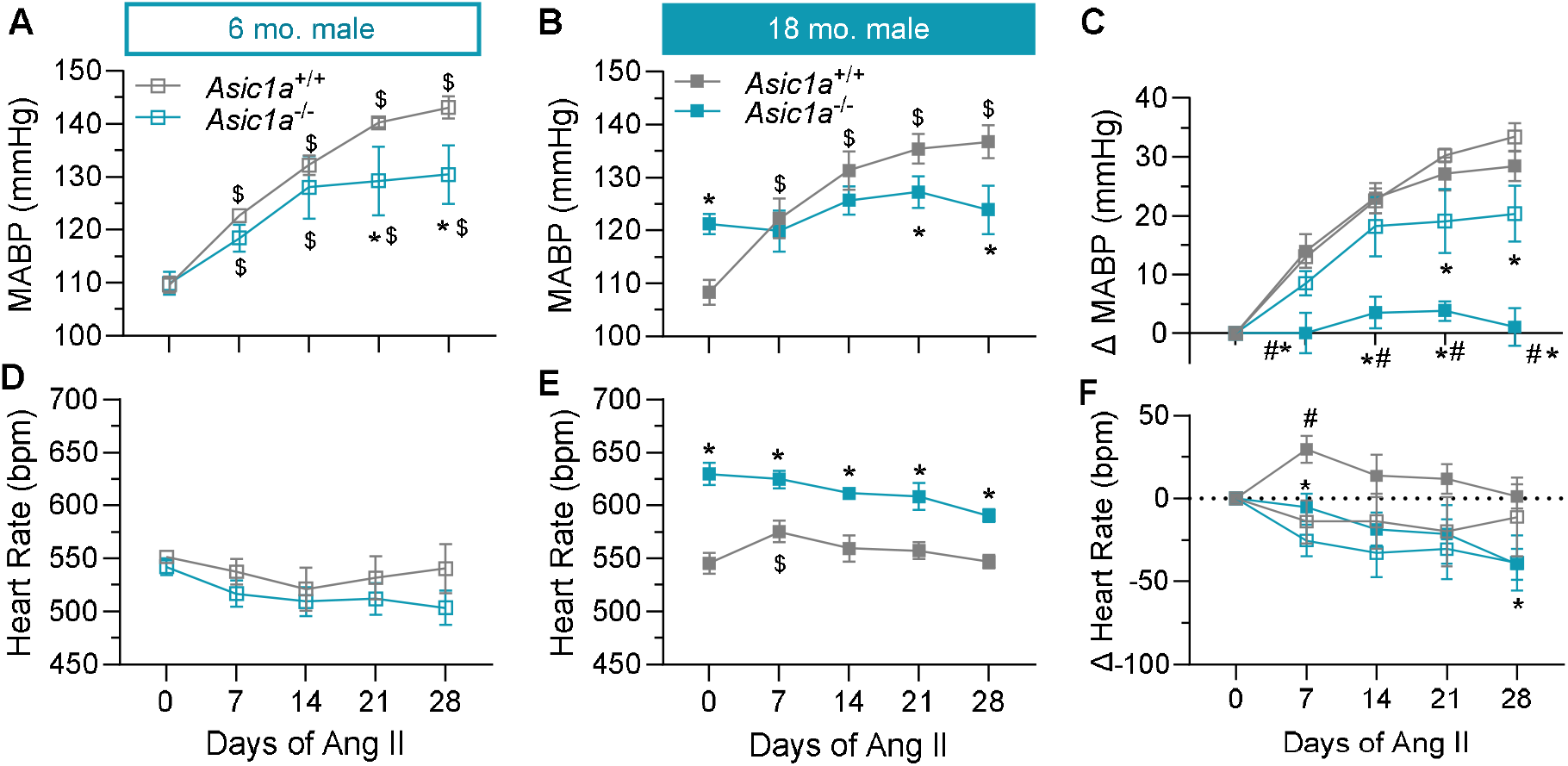
Asic1a deletion leads to age-dependent protection against Ang II-induced hypertension in male mice. **A,B)** MABP and **C)** changes in MABP in response to angiotensin II infusion (600ng/kg/min) over 28 days in **A)** adult (6-month-old) and **B)** aged (18-month-old) male mice. **D,E)** HR and **F)** changes in HR over the same time. n =5-6 animals per group; * p < 0.05 versus *Asic1a*^+/+^; $ p<0.05 vs time 0; # p<0.05 vs 6-month-old adult; analyzed by repeated measures two-way ANOVA, individual groups compared using Tukey’s multiple comparisons test.

Given our previous findings that smooth muscle-specific deletion of ASIC1a protects against hypoxia-induced pulmonary hypertension^11^, we examined its role in systemic Ang II-induced hypertension. However, unlike global deletion, smooth muscle ASIC1a deletion did not affect the development of Ang II-induced hypertension (**Figure S1**). Furthermore, global ASIC1a deletion had no impact on MABP or HR in response to Ang II in either young or aged female mice (**Figure S2**), demonstrating this sex-specific role of ASIC1a extends to Ang II-induced hypertension. While the resistance to Ang II in aged *Asic1a*^-/-^ males may be partially attributed to already elevated baseline MABP and HR, the blunted response observed in younger *Asic1a*^-/-^ mice suggests that additional mechanisms may be at play.

### Increased MABP and HR in aged male *Asic1a*^-/-^ mice are associated with elevated norepinephrine (NE)

Urine NE levels were significantly higher in aged male *Asic1a*^-/-^ mice (**Figure 3A**), with no differences observed in female mice (**Figure 3B**). Epinephrine (Epi) levels were not different across groups in either male or female mice (**Figure S3A-B**). These findings suggest that increased sympathetic outflow in aged male *Asic1a*^-/-^ mice contributes to neurogenic-based hypertension. Enhanced sympathetic nerve activity can arise from multiple causes^19^, and we have investigated several, including sensory afferent dysfunction, excess circulating hormones, hypernatremia, and chronic stress.

**Figure 3.**
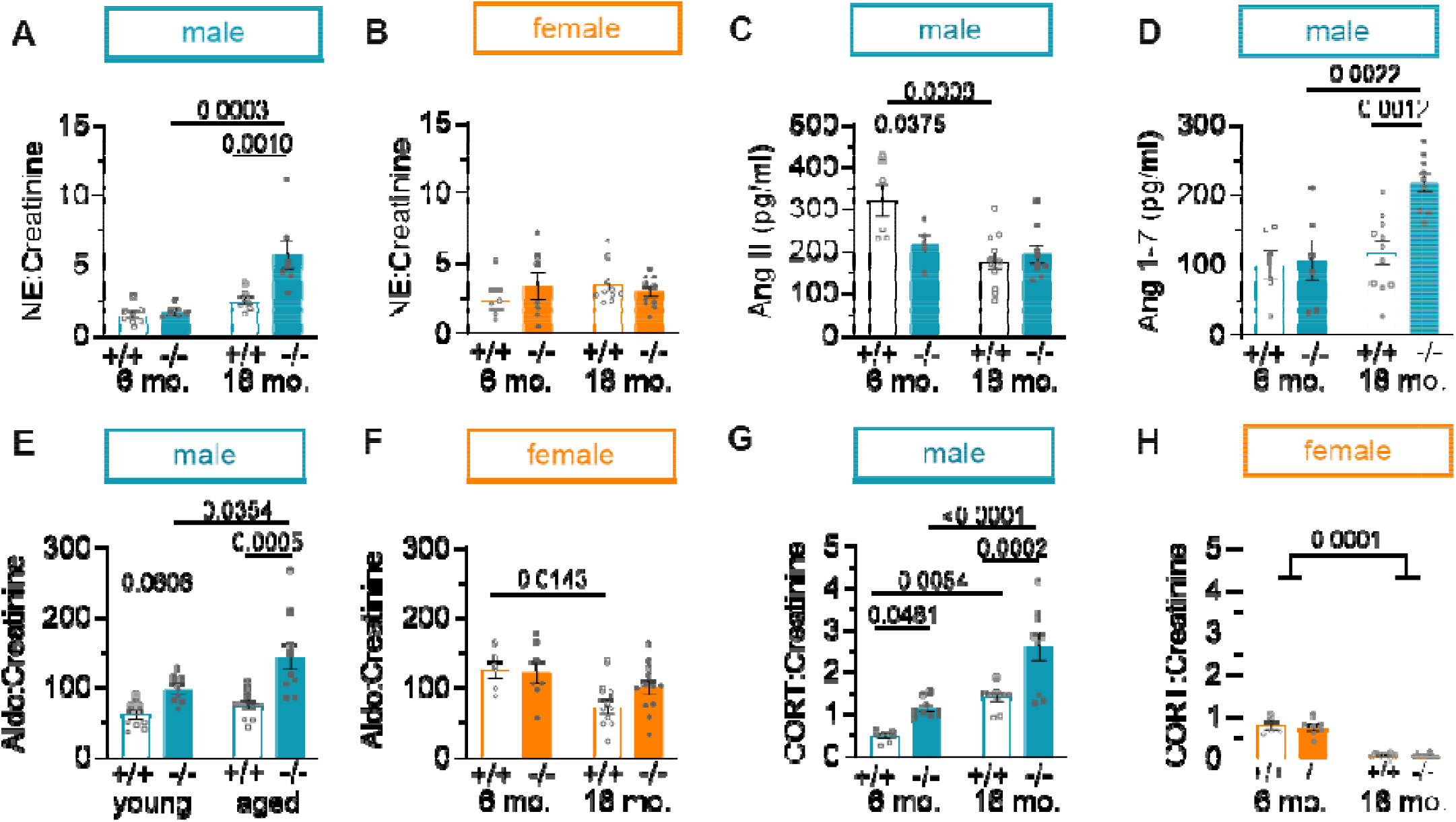
Norepinephrine, Aldosterone, and Cortisol are elevated in aged male *Asic1a*^-/-^ mice. Urine Norepinephrine (NE; ng/ml):Creatinine (mg/dl) ratio in **A**) male and **B**) female 6- and 18-month-old *Asic1a*^+/+^ and *Asic1a*^-/-^ mice. Plasma concentration of **C**) Angiotensin II (pg/ml) and **D**) Angiotensin 1-7 (pg/ml) in male mice. Urine aldosterone (pg/ml):Creatinine (mg/dl) ratio in **E**) male and **F)** female mice. Urine corticosterone (ng/ml): Creatinine (mg/dl) ratio in **G**) male and **H**) female mice. Analyzed by two-way ANOVA, individual groups were compared using Tukey’s multiple comparisons test; *n* = number of animals/group. Creatinine was not different between groups (**Figure S3C-D**).

Previous studies suggest that ASIC2 and ASIC3 contribute to sensory afferent activity^13,14^. Using whole-body plethysmography in conscious young mice, we previously demonstrated that the genetic deletion of ASIC1, 2, or 3 does not affect the hypercapnic ventilatory response in mice, though ASIC2 plays a minor role in the hypoxic ventilatory response^20^. Given the potential influence of aging and excess body mass in *Asic1a*^-/-^ mice, we examined ventilatory responses in aged animals. Aged male *Asic1a*^-/-^ mice exhibited a significantly greater increase in tidal volume in response to hypoxia compared to *Asic1a*^+/+^ mice, yet baseline minute ventilation and ventilatory responses to hypercapnia and hypoxia remained similar between groups (**Figure S4**).

### Aldosterone, independent of angiotensin II (Ang II), is associated with hypertension in aged male *Asic1a*^-/-^ mice

A significant increase in Ang II-mediated sympathetic nervous system activity is known to be the key driving force behind many types of hypertension^21^. However, Ang II levels were not elevated in aged male *Asic1a*^-/-^ mice. On the contrary, Ang II levels were lower in 6-month-old *Asic1a*^-/-^ compared to *Asic1a*^+/+^ male mice (**Figure 3C**). While there was no difference in Ang II levels in aged male mice, angiotensin 1-7 levels were significantly elevated (**Figure 3D**). Importantly, aldosterone levels were elevated in male *Asic1a*^-/-^ mice (**Figure 3E**), which was not observed in female mice (**Figure 3F**). These findings suggest that ASIC1a deficiency results in a renin-angiotensin-independent form of hyperaldosteronism which is often characterized by hypertension, hypokalemia, and alkalosis^22^. Consistent with this, we observe alkalosis with renal compensation in conscious, unrestrained male *Asic1a*^-/-^ mice as indicated by a significant reduction in arterial K^+^, pCO_2_, and HCO_3_^-^ (**Table S2**). There was no significant difference in Na^+^ or pH. Female mice exhibit alkalosis with an elevated pH compared to male mice, however K^+^ was not significantly different between groups.

### ASIC1a deficiency causes corticosterone excess in male, but not female mice

Aldosterone production is primarily regulated by Ang II, hyperkalemia, and adrenocorticotropic hormone (ACTH)^23^. Recent studies have identified cortisol (corticosterone in rodents) co-secretion as a key feature of primary aldosteronism, which may additionally contribute to cardiometabolic complications^24^. We found that both 6- and 18-month-old male *Asic1a*^-/-^ mice have elevated urine corticosterone levels (**Figure 3G**), a change that is absent in females (**Figure 3H**). To further investigate, we accessed publicly available RNA-Seq data (accession ID: GSE185718)^25^ from hypothalamic samples collected during the light and dark cycle from 3-month-old *Asic1a*^+/+^ and *Asic1a*^*-/-*^ mice. In *Asic1a*^+/+^ mice, corticotropin-releasing hormone (*Crh)* expression decreased during the active dark cycle relative to the resting light cycle, whereas in *Asic1a*^−/−^ mice, *Crh* remained elevated during the dark cycle (**Figure S5A**). This was accompanied by increased expression of *Hsd11b1* (11β-HSD1)—which activates corticosterone, and *Nr3c1—*the glucocorticoid receptor, in *Asic1a*^-/-^ mice during the dark cycle (**Figure S5B-C**). However, transcript levels of *Pomc* (ACTH precursor), *Fkbp5* (glucocorticoid co-chaperone), and *Nr3c2* (mineralocorticoid receptor) were unchanged (**Figure S5D-F**). Additionally, Peng et al. confirmed increased expression of *Fos* and *FosB (*neural activation markers), *Sgk1 (*serum glucocorticoid kinase 1, a corticosterone-responsive gene), and *Nr4a1* (a stress-responsive gene) during the dark cycle in *Asic1a*^-/-^ mice^25^. These data indicate that ASIC1a deficiency leads to early chronic stress through activation of the hypothalamus pituitary adrenal (HPA) axis that precedes the development of hypertension.

### Aged male *Asic1a*^-/-^ mice exhibit increased cardiac hypertrophy and aortic fibrosis

Prolonged exposure to elevated aldosterone and corticosterone can cause cardiac and renal damage independently of blood pressure^26^. While heart mass increased with age in both male and female mice, there was a further exacerbation of left ventricular mass in aged male *Asic1a*^-/-^ mice (**Figure 4A**). The individual cardiomyocyte cross-sectional area (CSA) also increased with age, and interestingly, both male and female *Asic1a*^-/-^ mice had significantly larger cardiomyocyte CSA regardless of age (**Figure 4B**), indicating a potential adaptive response to chronic cardiac stress in *Asic1a*^-/-^ mice independent of disease in both sexes. Both cardiac (**Figure S6A**) and coronary arterial fibrosis (**Figure S6B**) increased with age, independent of genotype. In contrast, aortic fibrosis correlated with changes in MABP and HR, with greater aortic fibrosis observed in aged female *Asic1a*^+/+^ and aged male *Asic1a*^-/-^ mice (**Figure 4C**).

**Figure 4.**
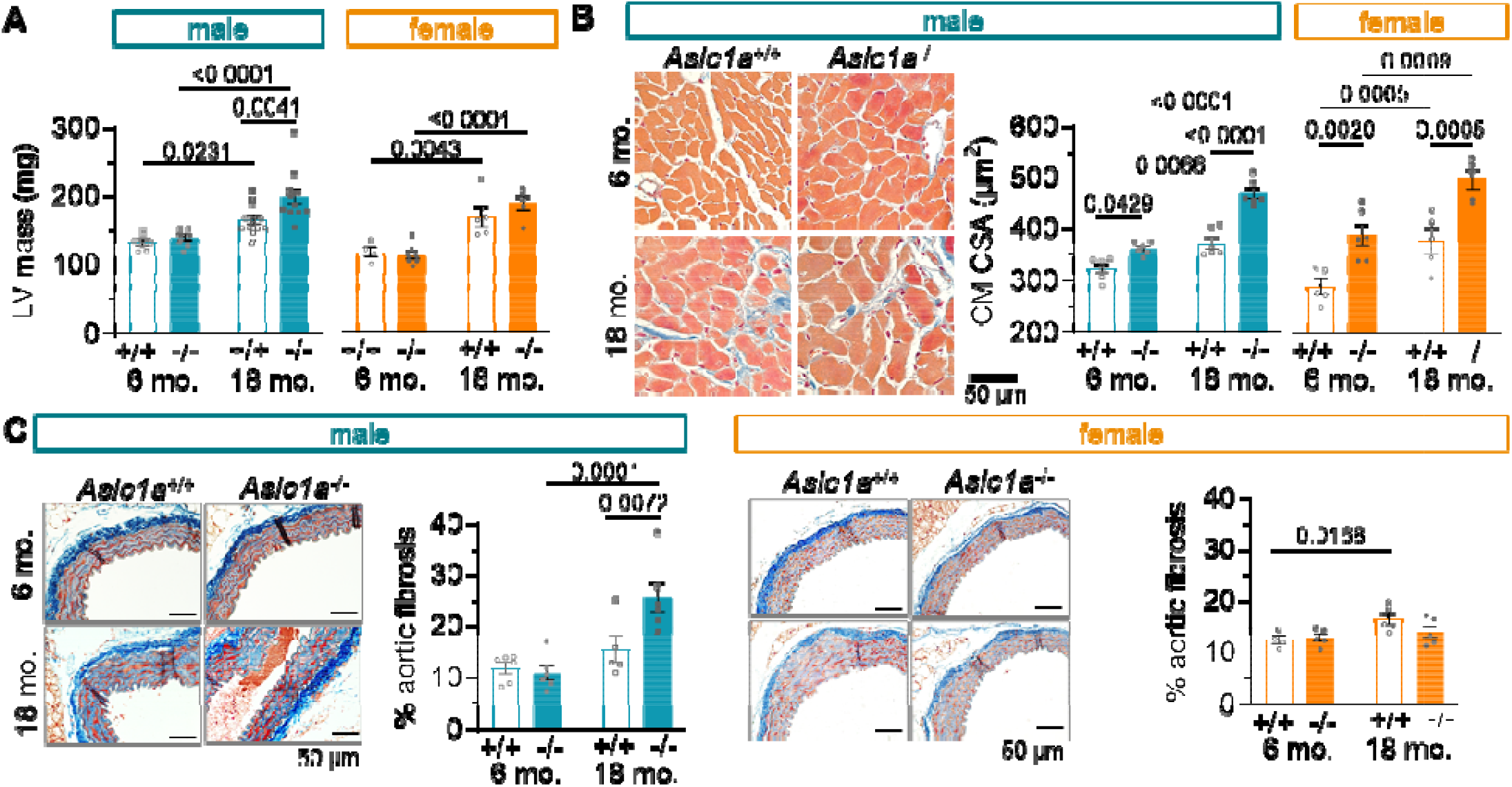
Aged male *Asic1a*^-/-^ mice exhibit increased cardiac hypertrophy and aortic fibrosis. **A**) Left ventricular mass (LV, mg). **B**) AZAN trichrome-stained whole heart sections (male only) and cardiomyocyte cross-sectional area (CM-CSA; μm^2^) from 6- and 18-month-old *Asic1a*^+/+^ and *Asic1a*^-/-^ mice. AZAN trichrome shows cell nuclei (dark red), collagen (blue), and orange-red in the cytoplasm. C) thoracic aorta fibrosis (% thresholded area). Values are mean ± SEM; n= animals/group; analyzed by two-way ANOVA, individual groups compared using Tukey’s multiple comparisons test.

### ASIC1a deficiency exacerbates glomerular remodeling and renal fibrosis in aged male mice

In male mice, aging significantly increased renal fibrosis and glomeruli CSA in both genotypes, but hypertensive aged male *Asic1a*^-/-^ mice exhibited a further increase in renal fibrosis (**Figure 5A-B**) and glomeruli CSA (**Figure 5C**), which was associated with a decrease in the number of glomeruli (**Figure 5D**). While renal fibrosis and glomeruli CSA are also increased in aged female mice, there is no significant difference between genotypes. However, aged female *Asic1a*^*+/+*^, but not *Asic1a*^*-/-*^, mice had fewer glomeruli (**Figure S7**). These findings link hypertension in aged male *Asic1a*^*−/−*^ mice to increased fibrosis and glomerular changes. To assess functional impact, we measured urine osmolality and protein concentration. In males, urine osmolality was lower in both 6- and 18-month-old *Asic1a*^*−/−*^ mice compared to *Asic1a*^*+/+*^ (**Figure 5E**). Together with the observation of lower hematocrit (**Table S2**) suggests volume expansion in male *Asic1a*^*−/−*^ mice which corresponds with the observed hypokalemia in these animals. While urine protein levels were elevated in aged mice, it was unaffected by genotype (**Figure 5F**). No differences in urine protein or osmolality were detected in females (**Figure S7E-F**).

**Figure 5.**
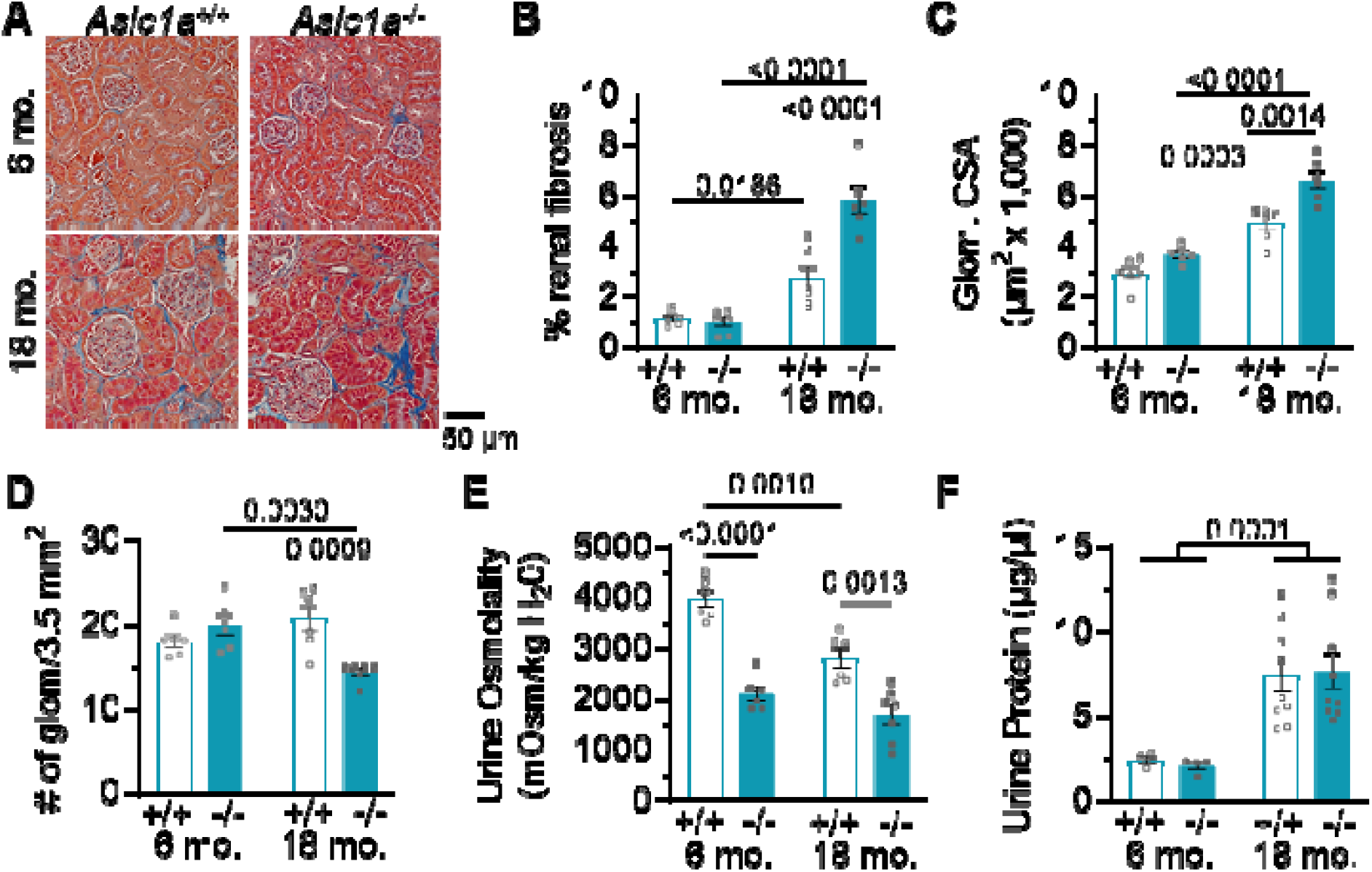
Asic1a deletion exacerbates glomerular remodeling and renal fibrosis in aged male mice. **A**) Representative AZAN trichrome-stained kidney sections and summary data showing **B**) percent renal fibrosis (% of 3.5 mm^2^ area) **C**) glomerular cross-sectional area (CSA; μm^2^ x 1,000), and **D**) glomeruli number (3.5 mm^2^ area). Measurements for urine **E**) osmolality (Osm/kg H2O) and **F**) protein (μg/μL) from 6- and 18-month-old *Asic1a*^+/+^ and *Asic1a*^-/-^ male mice. Values are mean ± SEM; analyzed by two-way ANOVA, individual groups compared using Tukey’s multiple comparisons test.

### Mineralocorticoid receptor inhibition reduces MABP and HR in aged male *Asic1a*^-/-^ mice

To determine the role of aldosterone in age-induced hypertension, aged male mice were treated with spironolactone, a mineralocorticoid receptor antagonist. Spironolactone significantly lowered MABP and HR in *Asic1a*^-/-^ mice, while having no effect in *Asic1a*^+/+^ controls, restoring MABP and HR in *Asic1a*^-/-^ mice to normotensive *Asic1a*^+/+^ mice (**Figure 6**). Additionally, spironolactone prevented the increase in NE, implicating mineralocorticoid receptor signaling in the elevated sympathetic activity. Together, these findings indicate that mineralocorticoid signaling plays a key role in the cardiovascular dysfunction associated with Asic1a deficiency.

**Figure 6.**
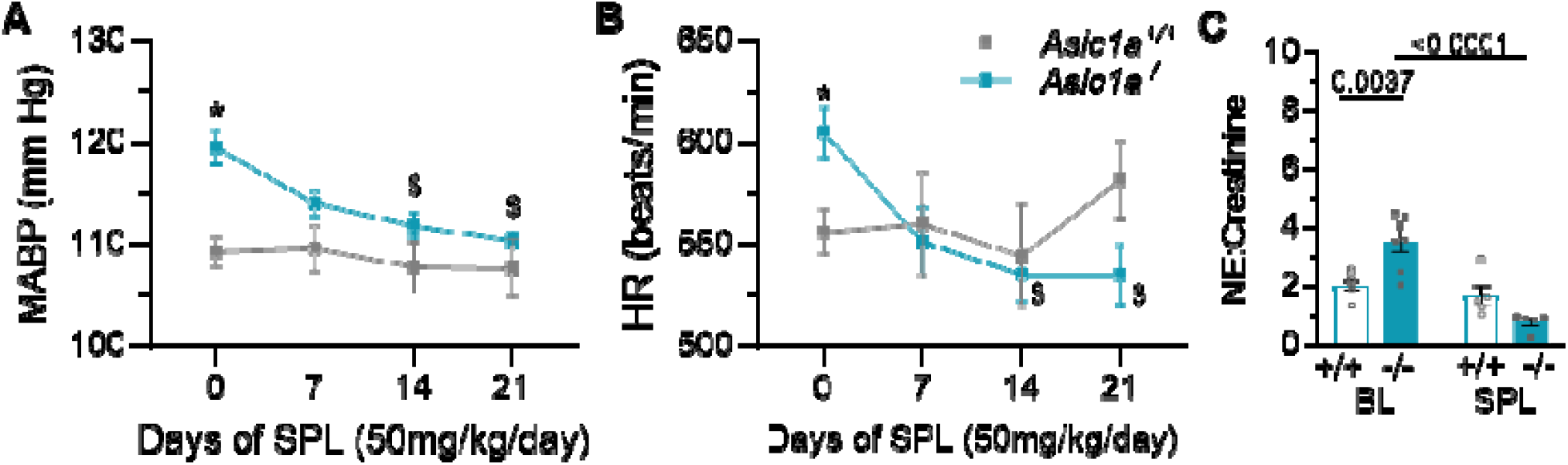
Inhibition of mineralocorticoid receptor lowers MABP and HR in aged *Asic1a*^-/-^ male mice. Twenty-four summary data showing **A**) MABP and **B**) HR in response to spironolactone (SPL; 50mg/kg/day) over 21 days in 18-month-old *Asic1a*^+/+^ and *ASIC1a*^-/-^ male mice. **C**) Urine Norepinephrine (NE; ng/ml):Creatinine (mg/dl) ratio at baseline (BL) and following 21 days SPL treatment. n =5-6 animals per group; * p < 0.05 versus *Asic1a*^+/+^; $ p<0.05 vs day 0; analyzed by repeated measures two-way ANOVA, individual groups

## DISCUSSION

This study investigates the role of ASIC1a in cardiovascular health and disease, revealing profound differences in ASIC1a’s regulation of cardiovascular function during various physiological states, namely sex, age, and Ang II-induced hypertension. We found that aged male *Asic1a*^-/-^ mice develop hypertension, driven by early chronic stress and subsequent aldosterone excess and activation of sympathetic nerve activity, whereas female *Asic1a*^-/-^ mice remain unaffected. Notably, hyperaldosteronism in male *Asic1a*^-/-^ mice occurs independent of the renin-angiotensin system and is associated with protection against Ang II-induced hypertension. These findings highlight the critical, sex-specific role of ASIC1a in cardiovascular regulation, where its deficiency in male mice triggers early corticosterone excess, leading to hyperaldosteronism and identifying ASIC1a deficiency as a potential mediator of endocrine-related hypertension.

The production of aldosterone is primarily regulated by Ang II, hyperkalemia, and ACTH^23^. In male *Asic1a*^-/-^ mice, the presence of aldosterone excess, hypokalemia, and volume expansion— in the absence of increased Ang II—suggests a renin-angiotensin system (RAS)-independent form of hyperaldosteronism, akin to primary aldosteronism^27,28^. The sustained *Crh* and elevated corticosterone levels following ASIC1a deficiency indicate dysregulation of the HPA axis may contribute to hyperaldosteronism. Given ASIC1a’s role in neurotransmission and synaptic plasticity, ASIC1a deletion may influence neuroendocrine function. Indeed, Peng et al. recently showed that signaling in the hypothalamus and pituitary are altered in male *Asic1a*^-/-^ mice leading to lower thyroid-releasing hormone (*Trh*) and body temperature during the dark cycle^25^. Using these data (accession ID: GSE185718)^25^, we further show other alterations in the hypothalamus leading to elevated dark cycle corticosterone signaling and a state of “chronic stress” that precedes hypertension. However, ASIC1a is also expressed in the kidney (proximal tubule, endothelial, and fibroblasts) and the adrenal gland (Schwann, smooth muscle, endothelial, and fibroblasts)^9^. Therefore, we cannot rule out the possibility that intra-adrenal or intrarenal ASIC1a deficiency affects corticosterone or aldosterone synthesis. Further studies are needed to delineate the potential link between ASIC1a deficiency, HPA axis dysregulation, chronic stress, and RAS-independent aldosterone excess.

RAS-independent aldosteronism is marked by suppression of Ang II signaling^28^, which may mediate resistance to Ang II-induced hypertension in male *Asic1a*^-/-^ mice. Interestingly, aged male *Asic1a*^-/-^ mice also exhibited a significant increase in Ang 1-7 levels—a peptide known to counteract Ang II signaling—which could also mediate the resistance to Ang II hypertension. This elevation may represent a compensatory response, as Ang 1-7 is an alternative cleave product of angiotensin I that has been shown to negatively regulate aldosterone^29,30^. These findings suggest that enhanced Ang 1-7 signaling in *Asic1a*^-/-^ mice could play a protective role in hyperaldosteronism by modulating blood pressure.

Elevated aldosterone and corticosterone are associated with a higher risk of cardiovascular complications, as both can cause cardiac and renal damage independent of blood pressure^31^. Since corticosterone binds mineralocorticoid receptors with equal affinity as aldosterone, elevated corticosterone levels can mimic aldosterone’s effects while contributing to metabolic complications through glucocorticoid receptors^32^. We observed many factors associated with hypertension—such as increased left ventricular mass, aortic fibrosis, renal fibrosis, and glomerular hypertrophy—that cannot be solely attributed to corticosterone, aldosterone, sympathetic activity, or blood pressure. However, we also identified several early structural and functional changes in male *Asic1a*^-/-^ mice that precede hypertension, including hypokalemia, reduced urine osmolality, increased pulse pressure (arterial stiffness), and cardiomyocyte hypertrophy. While corticosterone-binding mineralocorticoid receptors likely contribute to these effects, the presence of cardiomyocyte hypertrophy (indicated by CSA) in female *Asic1a*^-/-^ mice—despite unchanged corticosterone levels—suggests that ASIC1a deficiency may drive cardiomyocyte hypertrophy through alternative mechanisms.

Estrogen generally protects against cardiovascular disease in women, but this protection declines after menopause when estrogen levels decrease^33^. Acyclic female rodents do not exhibit as large of a decline in estrogen that resembles human menopause, however aged female mice reportedly have lower estrogen and higher progesterone levels^34^. While our data show that aged female *Asic1a*^-/-^ mice are protected from hypertension—seemingly contradicting this view—it aligns with sex-specific regulation of the HPA axis and corticosterone signaling. Estrogens enhance HPA axis activity and increase adrenal response to ACTH, whereas androgens suppress HPA activity^35–37^. Consequently, females typically have higher total circulating corticosterone levels. However, since 80-90% of circulating corticosterone is bound to corticosteroid-binding globulin (CBG)^38^—which is twice as high in females^39^—biologically active corticosterone is either similar between sexes or lower in females, limiting its bioavailability for negative feedback regulation. Additionally, females express lower levels of glucocorticoid and mineralocorticoid receptors, resulting in less efficient HPA axis negative feedback^35,40,41^. Notably, a recent report shows that hypothalamus-specific glucocorticoid receptor deletion disrupts feedback control in males but not in female mice, suggesting sex-specific differences in HPA axis regulation^41^.

Sex differences also extend to corticosterone metabolism. 11β-HSD1 converts inactive cortisone to active corticosterone, and 11β-HSD2 converts corticosterone to cortisone. Males exhibit a higher 11β-hydroxysteroid dehydrogenase types 1 and 2 (11β-HSD1/2) ratio, enhancing corticosterone signaling^42–44^. Furthermore, males exhibit enhanced corticosterone-mediated signaling via serum and glucocorticoid kinase 1 (SGK1)^45^, a key regulator of metabolic and cardiovascular pathways, highlighting the potential impact of sex-specific corticosterone regulation on disease susceptibility.

With aging, the loss of sex hormones results in greater HPA axis activation in older men compared to women, correlating with an elevated risk of cardiometabolic disease in men^46^. Although aging female rodents do not experience the same decline in estrogen that resembles human menopause, acyclic female mice exhibit lower estrogen and higher progesterone levels^34^. Consistent with this, we observed significantly lower corticosterone in aged female mice. These findings emphasize the critical role of sex-specific corticosterone regulation in shaping cardiometabolic disease risk with aging and implicate ASIC1a as a potential mediator.

## PERSPECTIVES

ASIC1a polymorphisms in humans have been linked to neuropathologies such as epilepsy and panic disorder^47,48^—both associated with elevated cortisol levels^49,50^—as well as cardiac and cerebrovascular ischemic injury^10^. Although the functional consequences of these polymorphisms regarding gain or loss of function remain unclear, *Asic1a*^-/-^ mice exhibit greater anxiety-like behaviors^51^ and increased epileptic seizure susceptibility^52^. Our findings suggest an additional crucial role for ASIC1a in neuroendocrine regulation and cardiovascular health, revealing striking sex differences in its function. In male mice, ASIC1a deficiency induces early corticosterone excess and subsequent aldosterone-driven hypertension. Given the association between chronic stress, hyperaldosteronism, diabetes, and resistant hypertension^53,54^— particularly in men^55^—future studies will investigate the mechanisms linking ASIC1a deficiency, endocrine dysregulation, and metabolic consequences. Additionally, a deeper understanding of ASIC1a polymorphisms and their genetic associations may facilitate earlier detection of cardiometabolic diseases.

## NOVELTY AND RELEVANCE

### What Is New?

- Identifies ASIC1a as a novel sex-specific regulator of cardiovascular function.
- Links ASIC1a deficiency in male mice to disrupted hypothalamus-pituitary-adrenal (HPA) axis regulation, leading to chronic stress that precedes hyperaldosteronism.

### What Is Relevant?

- Demonstrates that ASIC1a deficiency contributes to hypertension through hormonal dysregulation and sympathetic overactivity.
- Highlights sex differences in hypertension risk, particularly in aging males.
- Suggests ASIC1a as a potential target for understanding and treating endocrine-related hypertension.

### Clinical/Pathophysiological Implications

- ASIC1a polymorphisms may contribute to stress-related hypertension and metabolic disorders.
- Understanding ASIC1a’s role may lead to earlier detection of cardiometabolic disease and could inform personalized treatments.

## LIST OF ABBREVIATIONS (alphabetical)

ACTH: adrenocorticotropic hormone
Ang II: angiotensin II
ASIC1a (gene name ACCN2): acid-sensing ion channel 1a
CBG: corticosteroid-binding globulin
CSA: cross-sectional area
Epi: epinephrine (adrenaline)
HR: heart rate
HPA: hypothalamus-pituitary-adrenal
11β-HSD1/2: 11β-hydroxysteroid dehydrogenase types 1 and 2
MABP: mean arterial blood pressure
NE: norepinephrine (noradrenaline)
RAS: renin-angiotensin system
SGK1: serum and glucocorticoid kinase 1
SPL: spironolactone
SEM: standard error of the mean

## ACKNOWLEDGEMENTS

The authors thank Tamara Howard for assistance with tissue processing for immunohistochemistry and Lindsay Herbert and Sana Gul for their technical support in animal care and genotyping.

## SUPPLEMENTAL MATERIAL

### Data Supplement

Expanded Methods

Major Resources

Table Tables S1-S2

Figure S1-S7

References #20, 25, 56-–61

## REFERENCES

1. Waldmann R, Champigny G, Bassilana F, Heurteaux C, Lazdunski M. A proton-gated cation channel involved in acid-sensing. Nature. 1997;386:173–177.

2. Boscardin E, Alijevic O, Hummler E, Frateschi S, Kellenberger S. The function and regulation of acid-sensing ion channels (ASICs) and the epithelial Na+ channel (ENaC): IUPHAR Review 19. Br. J. Pharmacol. 2016;173:2671–2701.

3. Hoagland E, Sherwood T, Lee K, Walker C, Askwith C. Identification of a calcium permeable human acid-sensing ion channel 1 transcript variant. J Biol Chem. 2011;285:41852–41862.

4. Sherwood TW, Lee KG, Gormley MG, Askwith CC. Heteromeric acid-sensing ion channels (ASICs) composed of ASIC2b and ASIC1a display novel channel properties and contribute to acidosis-induced neuronal death. J Neurosci. 2011;31:9723–9734.

5. Xiong ZG, Zhu XM, Chu XP, Minami M, Hey J, Wei WL, MacDonald JF, Wemmie JA, Price MP, Welsh MJ, et al. Neuroprotection in ischemia: blocking calcium-permeable acid-sensing ion channels. Cell. 2004;118:687–698.

6. Yermolaieva O, Leonard AS, Schnizler MK, Abboud FM, Welsh MJ. Extracellular acidosis increases neuronal cell calcium by activating acid-sensing ion channel 1a. PNAS. 2004;101:6752–6757.

7. Wemmie J, Chen J, Askwith C, Hruska-Hageman A, Price M, Nolan B, Yoder P, Lamani E, Hoshi T, Freeman J, et al. The acid-activated ion channel ASIC contributes to synaptic plasticity, learning, and memory. Neuron. 2002;34:463–477.

8. Benarroch EE. Acid-sensing cation channels: structure, function, and pathophysiologic implications. Neurology. 2014;82:628–35.

9. Tissue expression of ASIC1 - Summary - The Human Protein Atlas [Internet]. [cited 2021 Jul 5];Available from: https://www.proteinatlas.org/ENSG00000110881-ASIC1/tissue

10. Redd MA, Scheuer SE, Saez NJ, Yoshikawa Y, Chiu HS, Gao L, Hicks M, Villanueva JE, Joshi Y, Chow CY, et al. Therapeutic Inhibition of Acid-Sensing Ion Channel 1a Recovers Heart Function After Ischemia-Reperfusion Injury. Circulation. 2021;144:947–960.

11. Garcia SM, Yellowhair TR, Detweiler ND, Ahmadian R, Herbert LM, Gonzalez Bosc LV, Resta TC, Jernigan NL. Smooth muscle Acid-sensing ion channel 1a as a therapeutic target to reverse hypoxic pulmonary hypertension. Front Mol Biosci. 2022;9:DOI: 10.3389/fmolb.2022.989809.

12. Garcia SM, Naik JS, Resta TC, Jernigan NL. Acid-sensing ion channel 1a activates IKCa/SKCa channels and contributes to endothelium-dependent dilation. J Gen Physiol. 2023;155:e202213173.

13. Lu Y, Ma X, Sabharwal R, Snitsarev V, Morgan D, Rahmouni K, Drummond HA, Whiteis CA, Costa V, Price M, et al. The Ion Channel ASIC2 is Required for Baroreceptor and Autonomic Control of the Circulation. Neuron. 2009;64:885–897.

14. Tan Z-Y, Lu Y, Whiteis CA, Simms AE, Paton JFR, Chapleau MW, Abboud FM. Chemoreceptor Hypersensitivity, Sympathetic Excitation, and Overexpression of ASIC and TASK Channels Before the Onset of Hypertension in SHR. Circ Res. 2010;106:536–545.

15. Cheng C-F, Kuo TBJ, Chen W-N, Lin C-C, Chen C-C. Abnormal Cardiac Autonomic Regulation in Mice Lacking ASIC3. BioMed Res. Int. 2014;2014:709159.

16. Faraci FM, Taugher RJ, Lynch C, Fan R, Gupta S, Wemmie JA. Acid-Sensing Ion Channels. Circ. Res. 2019;125:907–920.

17. Drummond HA, Xiang L, Chade AR, Hester R. Enhanced maximal exercise capacity, vasodilation to electrical muscle contraction, and hind limb vascular density in ASIC1a null mice. Physiol. Rep. 2017;5:e13368.

18. Nitta CH, Osmond DA, Herbert LM, Beasley BF, Resta TC, Walker BR, Jernigan NL. Role of ASIC1 in the development of chronic hypoxia-induced pulmonary hypertension. Am J Physiol Heart Circ Physiol. 2014;306:H41–52.

19. Guyenet PG, Stornetta RL, Souza GMPR, Abbott SBG, Brooks VL. Neuronal Networks in Hypertension. Hypertension. 2020;76:300–311.

20. Detweiler ND, Vigil KG, Resta TC, Walker BR, Jernigan NL. Role of acid-sensing ion channels in hypoxia- and hypercapnia-induced ventilatory responses. PLoS One. 2018;13:e0192724.

21. Lamptey RNL, Sun C, Layek B, Singh J. Neurogenic Hypertension, the Blood–Brain Barrier, and the Potential Role of Targeted Nanotherapeutics. Int. J. Mol. Sci. 2023;24:2213.

22. Conn JW. Primary aldosteronism. J. Lab. Clin. Med. 1955;45:661–664.

23. Hattangady N, Olala L, Bollag WB, Rainey WE. Acute and Chronic Regulation of Aldosterone Production. Mol. Cell. Endocrinol. 2012;350:151–162.

24. Arlt W, Lang K, Sitch AJ, Dietz AS, Rhayem Y, Bancos I, Feuchtinger A, Chortis V, Gilligan LC, Ludwig P, et al. Steroid metabolome analysis reveals prevalent glucocorticoid excess in primary aldosteronism. JCI Insight. 2017;2:e93136. 93136.

25. Peng Z, Ziros PG, Martini T, Liao X-H, Stoop R, Refetoff S, Albrecht U, Sykiotis GP, Kellenberger S. ASIC1a affects hypothalamic signaling and regulates the daily rhythm of body temperature in mice. Commun. Biol. 2023;6:1–13.

26. Savard S, Amar L, Plouin P-F, Steichen O. Cardiovascular Complications Associated With Primary Aldosteronism. Hypertension. 2013;62:331–336.

27. Byrd JB, Turcu AF, Auchus RJ. Primary Aldosteronism. Circulation. 2018;138:823–835.

28. Vaidya A, Mulatero P, Baudrand R, Adler GK. The Expanding Spectrum of Primary Aldosteronism: Implications for Diagnosis, Pathogenesis, and Treatment. Endocr. Rev. 2018;39:1057–1088.

29. Shefer G, Marcus Y, Knoll E, Dolkart O, Foichtwanger S, Nevo N, Limor R, Stern N. Angiotensin 1–7 Is a Negative Modulator of Aldosterone Secretion In Vitro and In Vivo. Hypertens. Dallas Tex 1979. 2016;68:378–384.

30. Xue B, Zhang Z, Johnson RF, Guo F, Hay M, Johnson AK. Central endogenous angiotensin-(1–7) protects against aldosterone/NaCl-induced hypertension in female rats. Am. J. Physiol. - Heart Circ. Physiol. 2013;305:H699–H705.

31. Otsuka H, Abe M, Kobayashi H. The Effect of Aldosterone on Cardiorenal and Metabolic Systems. Int. J. Mol. Sci. 2023;24:5370.

32. Gomez-Sanchez E, Gomez-Sanchez CE. The Multifaceted Mineralocorticoid Receptor. Compr. Physiol. 2014;4:965–994.

33. Barsha G, Mirabito Colafella KM, Walton SL, Gaspari TA, Spizzo I, Pinar AA, Hilliard Krause LM, Widdop RE, Samuel CS, Denton KM. In Aged Females, the Enhanced Pressor Response to Angiotensin II Is Attenuated By Estrogen Replacement via an Angiotensin Type 2 Receptor-Mediated Mechanism. Hypertension. 2021;78:128–137.

34. Nilsson ME, Vandenput L, Tivesten Å, Norlén A-K, Lagerquist MK, Windahl SH, Börjesson AE, Farman HH, Poutanen M, Benrick A, et al. Measurement of a Comprehensive Sex Steroid Profile in Rodent Serum by High-Sensitive Gas Chromatography-Tandem Mass Spectrometry. Endocrinology. 2015;156:2492–2502.

35. Panagiotakopoulos L, Neigh GN. Development of the HPA axis: Where and when do sex differences manifest? Front. Neuroendocrinol. 2014;35:285–302.

36. Sheng JA, Bales NJ, Myers SA, Bautista AI, Roueinfar M, Hale TM, Handa RJ. The Hypothalamic-Pituitary-Adrenal Axis: Development, Programming Actions of Hormones, and Maternal-Fetal Interactions. Front. Behav. Neurosci. [Internet]. 2021 [cited 2024 Jan 9];14. Available from: doi: 10.3389/fnbeh.2020.601939

37. Kokras N, Hodes GE, Bangasser DA, Dalla C. Sex differences in the hypothalamic– pituitary–adrenal axis: An obstacle to antidepressant drug development? Br. J. Pharmacol. 2019;176:4090–4106.

38. Lewis JG, Bagley CJ, Elder PA, Bachmann AW, Torpy DJ. Plasma free cortisol fraction reflects levels of functioning corticosteroid-binding globulin. Clin. Chim. Acta. 2005;359:189–194.

39. McCormick CM, Linkroum W, Sallinen BJ, Miller NW. Peripheral and Central Sex Steroids Have Differential Effects on the HPA Axis of Male and Female Rats. Stress. 2002;5:235– 247.

40. Pooley AE, Benjamin RC, Sreedhar S, Eagle AL, Robison AJ, Mazei-Robison MS, Breedlove SM, Jordan CL. Sex differences in the traumatic stress response: the role of adult gonadal hormones. Biol. Sex Differ. 2018;9:32.

41. Solomon MB, Loftspring M, de Kloet AD, Ghosal S, Jankord R, Flak JN, Wulsin AC, Krause EG, Zhang R, Rice T, et al. Neuroendocrine Function After Hypothalamic Depletion of Glucocorticoid Receptors in Male and Female Mice. Endocrinology. 2015;156:2843– 2853.

42. Belanger K, Sullivan JC. Sex Differences Of Renal11β-hsd1 And 11β-hsd2 Expression In A Doca-salt Model Of Hypertension. Hypertension. 2021;78:A45–A45.

43. DeSchoolmeester J, Palming J, Persson T, Pereira MJ, Wallerstedt E, Brown H, Gill D, Renström F, Lundgren M, Svensson MK, et al. Differences between men and women in the regulation of adipose 11β-HSD1 and in its association with adiposity and insulin resistance. Diabetes Obes. Metab. 2013;15:1056–1060.

44. Vierhapper H, Heinze G, Nowotny P. Sex-specific Difference in the Interconversion of Cortisol and Cortisone in Men and Women. Obesity. 2007;15:820–824.

45. Rusai K, Prókai A, Szebeni B, Mészáros K, Fekete A, Szalay B, Vannay Á, Degrell P, Müller V, Tulassay T, et al. Gender differences in serum and glucocorticoid regulated kinase-1 (SGK-1) expression during renal ischemia/reperfusion injury. Cell. Physiol. Biochem. Int. J. Exp. Cell. Physiol. Biochem. Pharmacol. 2011;27:727–738.

46. Traustadóttir T, Bosch PR, Matt KS. Gender differences in cardiovascular and hypothalamic-pituitary-adrenal axis responses to psychological stress in healthy older adult men and women. Stress Amst. Neth. 2003;6:133–140.

47. Lv R-J, He J-S, Fu Y-H, Zhang Y-Q, Shao X-Q, Wu L-W, Lu Q, Jin L-R, Liu H. ASIC1a polymorphism is associated with temporal lobe epilepsy. Epilepsy Res. 2011;96:74–80.

48. Gugliandolo A, Gangemi C, Caccamo D, Currò M, Pandolfo G, Quattrone D, Crucitti M, Zoccali RA, Bruno A, Muscatello MRA. The RS685012 Polymorphism of ACCN2, the Human Ortholog of Murine Acid-Sensing Ion Channel (ASIC1) Gene, is Highly Represented in Patients with Panic Disorder. NeuroMolecular Med. 2016;18:91–98.

49. Cano-López I, González-Bono E. Cortisol levels and seizures in adults with epilepsy: A systematic review. Neurosci. Biobehav. Rev. 2019;103:216–229.

50. Abelson JL, Khan S, Liberzon I, Young EA. HPA axis activity in patients with panic disorder: review and synthesis of four studies. Depress. Anxiety. 2007;24:66–76.

51. Pidoplichko VI, Aroniadou-Anderjaska V, Prager EM, Figueiredo TH, Almeida-Suhett CP, Miller SL, Braga MFM. ASIC1a activation enhances inhibition in the basolateral amygdala and reduces anxiety. J. Neurosci. Off. J. Soc. Neurosci. 2014;34:3130–3141.

52. Ziemann AE, Schnizler MK, Albert GW, Severson MA, Howard Iii MA, Welsh MJ, Wemmie JA. Seizure termination by acidosis depends on ASIC1a. Nat Neurosci. 2008;11:816–822.

53. Huby A-C, Antonova G, Groenendyk J, Gomez-Sanchez CE, Bollag WB, Filosa JA, de Chantemèle EJB. Adipocyte-Derived Hormone Leptin Is a Direct Regulator of Aldosterone Secretion, Which Promotes Endothelial Dysfunction and Cardiac Fibrosis. Circulation. 2015;132:2134–2145.

54. Adler GK, Murray GR, Turcu AF, Nian H, Yu C, Solorzano CC, Manning R, Peng D, Luther JM. Primary Aldosteronism Decreases Insulin Secretion and Increases Insulin Clearance in Humans. Hypertension. 2020;75:1251–1259.

55. Hatano Y, Sawayama N, Miyashita H, Kurashina T, Okada K, Takahashi M, Matsumoto M, Hoshide S, Sasaki T, Nagashima S, et al. Sex-specific Association of Primary Aldosteronism With Visceral Adiposity. J. Endocr. Soc. 2022;6:bvac098.

56. Snow JB, Norton CE, Sands MA, Weise-Cross L, Yan S, Herbert LM, Sheak JR, Gonzalez Bosc LV, Walker BR, Kanagy NL, et al. Intermittent Hypoxia Augments Pulmonary Vasoconstrictor Reactivity through PKCβ/Mitochondrial Oxidant Signaling. Am. J. Respir. Cell Mol. Biol. 2020;62:732–746.

57. Sturgis LC, Cannon JG, Schreihofer DA, Brands MW. The role of aldosterone in mediating the dependence of angiotensin hypertension on IL-6. Am. J. Physiol.-Regul. Integr. Comp. Physiol. 2009;297:R1742–R1748.

58. Kiernan JA. In: Histological and Histochemical Methods: Theory and Practice. Scion Publishing; 2008. p. 205–207.

59. Ruifrok AC, Johnston DA. Quantification of histochemical staining by color deconvolution. Anal. Quant. Cytol. Histol. 2001;23:291–299.

60. Wu P-Y, Huang Y-Y, Chen C-C, Hsu T-T, Lin Y-C, Weng J-Y, Chien T-C, Cheng IH, Lien C-C. Acid-Sensing Ion Channel-1a Is Not Required for Normal Hippocampal LTP and Spatial Memory. J. Neurosci. 2013;33:1828–1832.

61. Wirth A, Benyó Z, Lukasova M, Leutgeb B, Wettschureck N, Gorbey S, Orsy P, Horváth B, Maser-Gluth C, Greiner E, et al. G12-G13-LARG-mediated signaling in vascular smooth muscle is required for salt-induced hypertension. Nat. Med. 2008;14:64–68.

