## Supplemental data for "Acid-Sensing Ion Channel 1a Deficiency Drives Endocrine Hypertension in Male Mice"

**EXPANDED METHODS**

**Animal Ethics**

Animals were housed 1-5 per cage in a specific pathogen-free animal care facility and maintained on a reverse 12:12-h light-dark cycle. Standard chow (Teklad soy protein-free diet #2920) and water were provided *ad libitum*. Animals were randomly allocated to experimental groups, and when possible, genotype and treatment assignments were blinded to the investigators. All protocols used in this study abide by the National Institutes of Health guidelines for animals and were reviewed by the Institutional Animal Care and Use Committee of the University of New Mexico School of Medicine (Protocol #22-201292-HSC). All animals were anesthetized with a fatal dose of pentobarbital sodium (200mg/kg, i.p.) and immediately euthanized by exsanguination after the loss of consciousness.

**Transgenic Mouse Models**

Studies were completed in male and female, 6- and 18-month-old, wildtype (*Asic1a^+/+^*), global ASIC1a knockout (*Asic1a^-/-^*), and inducible smooth muscle-specific ASIC1a knockout (MHC^CreER^-*Asic1a* ^fl/fl^) mice, as shown in **Major Resource Table**. Transgenic mice that have been backcrossed on a C57BL/6J background for at least 10 generations. For global ASIC1a knockout, *Asic1a* disruption was assessed by PCR amplification of DNA extracted from tail biopsies using REDExtract-N_Amp PCR Ready Mix (Sigma-Aldrich, XNAT). The PCR product was separated using gel electrophoresis on a 3% agarose gel and stained with ethidium bromide for visualization under UV light.

To selectively induce the deletion of Asic1a in smooth muscle, floxed (Asic1a^fl/fl^) mice were crossed with smooth muscle-specific Cre (MHC^CreER^) transgenic mice. Male mice were studied exclusively since the expression of iCreER^T2^ under the control of the SM promoter is inserted on the Y chromosome. For induction of Cre activity, MHC^CreER^-*Asic1a* ^fl/fl^ mice were injected with 75 mg/kg tamoxifen in corn oil, once daily for five consecutive days. Following 14 days, Cre recombinase was assessed from tail DNA using the 5′ LoxP forward and 3′ LoxP reverse primer pair (**Major Resource Table**).

**Assessment of Systemic Mean Arterial Blood Pressure (MABP) and Heart Rate (HR).** MABP and HR were measured in mice using radiotelemetry devices [PA-C10 and HD-X10; Data Systems International (DSI)]. Telemetry transmitters were implanted under inhaled isoflurane anesthesia (2% isoflurane and 98% O_2_ gas mixture). Buprenorphine (0.1 mg/kg; IM) was administered pre-surgery for pain management and recovery. Using sterile techniques, a midline incision was made to expose the carotid artery by blunt dissection. A small incision in the carotid artery was created between two silk sutures to insert the telemeter, which was advanced toward the heart and secured in place. The transmitter body was positioned subcutaneously in the mid-flank, and the incision was closed with sterile sutures. After a 5-day recovery period, MABP and HR were measured using Ponemah® Software for either a 72-hour baseline recording (10 seconds every 15 minutes) or once a week for 24-hours with the following treatments. Animals that did not survive the surgery or telemeters stopped working during the experiment were excluded from the study.

Angiotensin II (Ang II)-induced hypertension. Osmotic minipumps (Alzet 1002) were implanted under sterile conditions with inhaled isoflurane anesthesia (2% isoflurane, 98% O_2_) in the upper flank of mice. Buprenex (0.1 mg/kg, IM) was given pre-surgery for pain management. The pumps delivered Ang II at 600 ng/kg/min for 28 days.

Mineralocorticoid receptor inhibition. Spironolactone (SPL; 50 mg/kg/day) was administered orally to aged male mice once daily for 21 days via a 1:1 paste of ground chow and peanut butter. We and others have used this method to administer drugs successfully^56,57^.

**Whole-body plethysmography**

Whole-body plethysmography was used to assess respiratory frequency, tidal volume, and minute ventilation in conscious, unrestrained mice as previously described^20^. The plethysmography chamber consists of a ~50 cm^3^ transparent polycarbonate cylinder fitted with a Validyne DP45-16 differential pressure transducer. A controlled leak, created using a 50 μL glass syringe with the plunger removed, served as a high-pass filter to stabilize pressure changes within the chamber. Tidal volume (V_T_) was calculated using the following equation and normalized to body mass:

$V_{T}= \frac{P_{T}}{P_{K}} \times V_{K}\times\frac{T_{R}\left( P_{B}-P_{C} \right)}{T_{R}\left( P_{B}-P_{C} \right)-T_{C}\left( P_{B}-P_{R} \right)}$ , where:

V_K_ = the volume of air injected into the animal chamber for calibration

P_T_ = the pressure deflection associated with each tidal volume

P_K_ = the pressure deflection associated with injection of the calibrating volume, V_K_

T_R_ = body temperature, assumed to be for all animals

T_C_ = the air temperature in the chamber, which varied from 20–23°C

P_B_ = barometric pressure, 630 mmHg in Albuquerque, NM

P_R_ = vapor pressure of water at body temperature

P_C_ = vapor pressure of water in the chamber, derived from T_C_ assuming 100% humidity which was confirmed in pilot experiments (gas mixtures were humidified by bubbling through three consecutive flasks of water prior to entry into the plethysmography chamber).

**Sample Collection**

Urine**.** Urine was collected between 7:00 and 9:00 AM, corresponding to the beginning of light (inactive) cycle. Mice were gently held over parafilm while applying pressure to lower back to stimulate urination. If a mouse did not urinate within a couple of minutes, it was returned to its cage, and collection was attempted on a different day to minimize stress-related alterations in urine composition. Urine measurements were normalized to creatinine levels, which were similar across groups (**Figure S3C-D**). Biological assays were performed using a microplate reader (Tecan Infinite® M Plex), Qubit Fluorometer (Protein assay), and Advanced Instruments Osmometer (osmolality) according to the manufacturer’s instructions (**Major Resource Table**).

Plasma. Following deep anesthetization, blood (approximately 1 ml) was collected via blind cardiac puncture using a 25-gauge needle and transferred to an EDTA vacutainer containing protease inhibitor cocktail [in mM: 0.5 1,10-phenanthroline monohydrate, 0.125 pepstatin, 1 Na p-hydroxymercuribenzoate, and 0.003 rat renin inhibitor]. Plasma was separated by centrifugation at 2,000 rpm for 10 minutes at 4°C, aliquoted, and frozen for subsequent analysis of Angiotensin II and Angiotensin 1-7 by the Biomarker Analytical Core at Wake Forest University School of Medicine. Any sample with noticeable hemolysis was excluded from the study.

Tissue**.** The heart, aorta, and kidneys were harvested. The left ventricle was weighed and the tissue was fixed in 2% paraformaldehyde and subsequently embedded in paraffin. Tissue sections (5 µm thick) were mounted on Superfrost™ Plus slides (Fisher Scientific) and stained with Heidenhain’s Azan trichrome stain^1^. To assess fibrosis, 5 random images were taken using Nikon Eclipse E400 upright microscope and either a 4x (cardiac and renal; 3.5 mm^2^ area) or 20x objective (coronary artery and aorta; 0.145 mm^2^ area). The Color Deconvolution plugin for Image J was used to separate the blue collagen immunohistochemical stain^59^. Images were thresholded and % blue stain per area was calculated. Regions of interest were drawn around individual cardiomyocytes and glomeruli to determine cross-section area (μm^2^; ~200 measurements/animal).

**Femoral artery catheterization and blood electrolyte measurements**. Six-month-old male mice were chronically instrumented with tapered polyethylene (PE-10) femoral artery catheters under isoflurane anesthesia (5% isoflurane for induction of anesthesia, ~2% for maintenance)^20^. Post-operatively, mice received buprenorphine (0.05–0.1 mg/kg, s.c.) and enrofloxacin (15 mg/kg, s.c.) to reduce pain and prevent infection. Catheters were filled with 100 units/ml of heparinized saline and routed outside the cage via spring tethers to allow blood sampling from conscious, unrestrained mice. Blood sampling occurred two days post-surgery, with ~100 μL of blood collected directly onto Abbott iStat handheld blood gas analyzer cartridges (EG6+, Abbott Park, IL) to measure Na^+^, K^+^, HCO3^-^, and base excess. We previously reported PO_2_, PCO_2_, and pH from this dataset^2^. Here, we reanalyzed these data to examine sex as a biological variable.

**Differential Gene Expression Analysis**

Publicly available gene expression data were obtained from the Gene Expression Omnibus (GEO) database (accession ID: GSE185718)^25^. This dataset includes hypothalamic samples collected from 3-month-old *Asic1a*^+/+^ and *Asic1a^-/-^* mice at Zeitgeber Time (ZT) 1 and ZT13, corresponding to 1 hr after the start of the light and dark cycle, respectively (*n* = 4 mice/group). According to Peng et al.^25^ library sizes were scaled using TMM normalization and log-transformed into counts per million (CPM, EdgeR package version 3.28.1). Processed transcripts per million (TPM) expression data were downloaded and used to identify differentially expressed genes (DEGs) between experimental groups.

**MAJOR RESOURCE TABLE**

| **Experimental Mice** | **Source** | | | **RRID** | **Ref** | | **Genotyping Primers 5’→3’** | | | | **bp** |
| --- | --- | --- | --- | --- | --- | --- | --- | --- | --- | --- | --- |
| Asic1a^+/+^  (C57BL/6J) | The Jackson Laboratory | | | IMSR_JAX:000664 |  | | ***Asic1a* ^+/+^** **forward**: CATGTCACCAAGCTCGACGAGGTG | | | | 262 |
| Asic1a ^−/−^  (B6.129-Asic1 ^tm1Wsh^/J) | The Jackson Laboratory | | | IMSR_JAX:013733 | 7 | | ***Asic1a* ^−/−^** **forward**: TGGATGTGGAATGTGTGCGA **^+/+^** **and** **^−/−^** **reverse**: CCGCCTTGAGCGGCAGGTTTAAAGG | | | | 310 |
| Asic1a fl/fl  (B6.129-Asic1 tm1Lien) | National Laboratory Animal Center | | | RMRC #: 13158 | 60 | | ***5′LoxP forward:*** TCCTCTCCCAAACACACAC ***5′LoxP reverse:*** GAGTTCCCTCCAGATGTGAG ***3′ LoxP forward:*** AGGCCTGCAAACTGTCATCT ***3′ LoxP reverse:*** GTTGCATCTTGAGCCTCCTC | | | | 410 ^fl/fl^  314 ^+/+^  406 ^fl/fl^  306 ^+/+^ |
| MHC^CreER^  (B6.FVB-Tg (Myh11-icre/ERT2)1Soff/J) | The Jackson Laboratory | | | IMSR_JAX:019079 | 61 | | ***Cre Forward:*** TGACCCCATCTCTTCACTCC ***Cre Reverse:*** AGTCCCTCACATCCTCAGGTT | | | | 287 |
| **Biological Assay** | | | ***Company*** | | | | | ***Cat #*** | | ***Sample*** | |
| Epinephrine/Norepinephrine ELISA | | | Abnova | | | | | KA1877 | | urine | |
| Creatinine Colorimetric Assay | | | Cayman Chemical | | | | | 500701 | | urine | |
| Angiotensin (II, 1-7) | | | Wake Forest University | | | | | Biomarker Analytical Core | | Plasma + inhibitor cocktail | |
| Aldosterone Parameter Assay | | | R&D Systems | | | | | KGE016 | | urine | |
| Corticosterone ELISA kit | | | R&D Systems | | | | | KGE009 | | urine | |
| Qubit Protein Assay Kit | | | Invitrogen | | | | | A50668 | | urine | |
| Urine Osmolality | | | Advanced Instruments | | | | | Model #3320 | | urine | |
| **Chemicals/Biologicals** | | ***Description*** | | | | ***Company*** | | | ***Cat#*** | ***Concentration*** | |
| Renin Inhibitor Peptide, WFML | | Renin inhibitor | | | | AnaSpec | | | AS-60463-1 | 0.003 mM | |
| 1,10-phenanthroline monohydrate (o-PT) | | Metalloprotease inhibitor | | | | Sigma-Aldrich | | | P-1294 | 0.5 mM | |
| pepstatin | | Acid protease inhibitor | | | | Peninsula Labs | | | 4036 | 0.125 mM | |
| Na p-hydroxymercuribenzoate (NaHMB) | | Protease inhibitor | | | | Sigma-Aldrich | | | H0642 | 1 mM | |
| Tamoxifen | | Activates Cre recombinase | | | | Sigma-Aldrich | | | 85256 | 75 mg/kg for 5 days | |
| Angiotensin II | | Ang II, human | | | | Sigma-Aldrich | | | A9525 | 600 ng/kg/min for 28 days | |
| Spironolactone | | Mineralocorticoid receptor antagonist | | | | Thermo Scientific | | | AC207460010 | 50 mg/kg/day for 21 days | |

**SUPPLEMENTAL TABLES**

**Table S1.** Average body mass and baseline cardiovascular measurements in adult (6-month-old) and aged (18-month-old) *Asic1a*^+/+^ and *Asic1a*^-/-^ female and male mice.

| **female** | **6 mo.** | | **18 mo.** | |
| --- | --- | --- | --- | --- |
|  | *Asic1a*^+/+^ (5) | *Asic1a*^-/-^ (6) | *Asic1a*^+/+^ (9) | *Asic1a*^-/-^ (8) |
| body mass (grams) | 30 ± 3 | 28 ± 1 | 38 ± 2 | **41 ± 4 ^#^** |
| Systolic BP (mm Hg) | 124 ± 2 | 126 ± 3 | 129 ± 2 | 128 ± 2 |
| Diastolic BP (mm Hg) | 93 ± 2 | 96 ± 5 | 99 ± 1 | 95 ± 2 |
| Pulse pressure | 30.9 ± 2 | 32.8 ± 3 | 32 ± 1 | 34 ± 2 |
| Inactive (light) MABP | 99 ± 2 | 105 ± 3 | **111 ± 2 ^#^** | 108 ± 2 |
| Inactive (light) HR | 563 ± 10 | 560 ± 8 | **620 ± 7 ^#^** | **587 ± 9 *** |
| Active (dark) MABP | 118 ±2 | 112 ± 2 | 117 ± 2 | 114 ± 2 |
| Active (dark) HR | 643 ± 18 | 587 ± 11 | 622 ± 16 | 607 ± 10 |
| **male** | **6 mo.** | | **18 mo.** | |
|  | *Asic1a*^+/+^ (13) | *Asic1a*^-/-^ (13) | *Asic1a*^+/+^ (9) | *Asic1a*^-/-^ (6) |
| body mass (grams) | 34 ± 1 | 33 ± 2 | 41 ± 1 | **51 ± 3 * ^#^** |
| Systolic BP (mm Hg) | 125 ± 3 | 129 ± 6 | 122 ± 8 | **136 ± 5 *** |
| Diastolic BP (mm Hg) | 99 ± 6 | 98 ± 6 | 97 ± 6 | **107 ± 6 * ^#^** |
| Pulse pressure | 26 ± 2 | **31 ± 2** * | 26 ± 2 | **29 ± 2 *** |
| Inactive (light) MABP | 105 ± 3 | 107 ± 5 | 105 ± 7 | **115 ± 5 * ^#^** |
| Inactive (light) HR | 524 ± 6 | 518 ± 11 | 548 ± 11 | **618 ± 12 * ^#^** |
| Active (dark) MABP | 119 ± 7 | 120 ± 6 | 114 ± 8 | **128 ± 5 *** |
| Active (dark) HR | 579 ± 11 | 583 ± 7 | 573 ± 16 | **644 ± 6 * ^#^** |

*Cardiovascular measurements were averaged over 72 hours. Values are mean ± SEM; n is indicated in parentheses as animals/group, analyzed by three-way ANOVA, significant interactions between the individual groups were compared using Tukey’s multiple comparisons test where * p<0.05 vs corresponding Asic1a^+/+^, # p < 0.05 vs corresponding 6 mo. old animal, and $ p < 0.05 vs corresponding female.*

**Table S2.** ***Asic1a*^-/-^ mice exhibit alkalosis with renal compensation.**

|  | **male** | | **female** | |
| --- | --- | --- | --- | --- |
|  | *Asic1a*^+/+^ (10) | *Asic1a*^-/-^ (5) | *Asic1a*^+/+^ (8) | *Asic1a*^-/-^ (5) |
| Na^+^, mmol/L | 152.2 ± 0.9 | 153.5 ± 1.7 | 153.4 ± 0.8 | 152.6 ± 1.3 |
| K^+^, mmol/L | 4.36 ± 0.05 | **4.06 ± 0.05 *** | 4.23 ± 0.07 | 4.18 ± 0.09 |
| pH | 7.36 ± 0.01 | 7.36 ± 0.01 | 7.38 ± 0.01 | **7.40 ± 0.01 ^$^** |
| pCO_2_, mm/Hg | 28.8 ± 0.4 | **26.4 ± 0.3 *** | 28.8 ± 0.9 | **25.7 ± 0.4 *** |
| pO_2_, mm/Hg | 61.9 ± 0.9 | **66.8 ± 0.8 *** | 60.8 ± 1.1 | **66.3 ± 0.9 *** |
| HCO_3_^-^, mmol/L | 16.88 ± 0.31 | **15.33 ± 0.34 *** | 17.34 ± 0.45 | **15.88 ± 0.21 *** |
| BE, mmol/L | -9.67 ± 0.56 | -10.00 ± 0.41 | **-8.13 ± 0.55 ^$^** | **-8.65 ± 0.24 ^$^** |
| Hct, % PCV | 44 ± 1 | **40 ± 4** * | **39 ± 2 ^$^** | 39 ± 2 |
| Hb, g/dL | 14.6 ± 0.2 | 14.1 ± 0.4 | 15.1 ± 0.1 | **14.1 ± 0.6 *** |

*Arterial blood gas measurements were taken from conscious, unrestrained ~4-month-old mice chronically instrumented with femoral artery catheters. Values are mean ± SEM; n is indicated in parentheses as animals/group, analyzed by two-way ANOVA, significant interactions between the individual groups were compared using* *Šídák's multiple comparisons test where * p<0.05 vs corresponding Asic1a^+/+^ and $ p < 0.05 vs corresponding female.BE: base excess, Hct: hematocrit, Hb: hemoglobin.*

**SUPPLEMENTAL FIGURES**

**Figure S1.** **Smooth Muscle Specific deletion of Asic1a does not affect baseline or Ang II-induced hypertension. A**) MABP and **B**) HR over 28 days of angiotensin II infusion (600ng/kg/min) in 6-month-old inducible smooth muscle-specific ASIC1a knockout mice (SM-*Asic1a*^-/-^ ), in the absence (-; grey) or presence (+; teal) of tamoxifen-induced gene recombination. n =5-6 animals per group; $ p<0.05 vs time 0; analyzed by repeated measures two-way ANOVA, individual groups compared using Tukey’s multiple comparisons test.

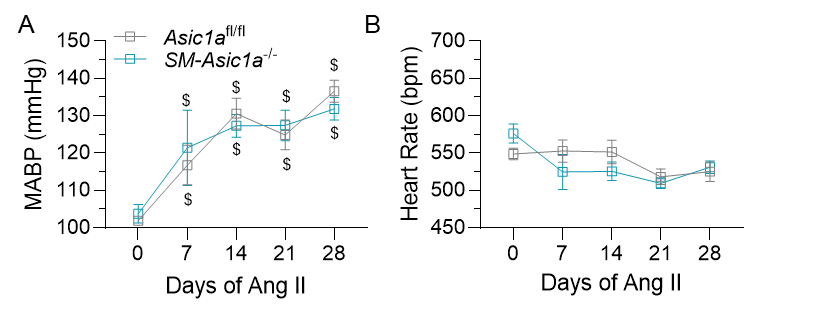

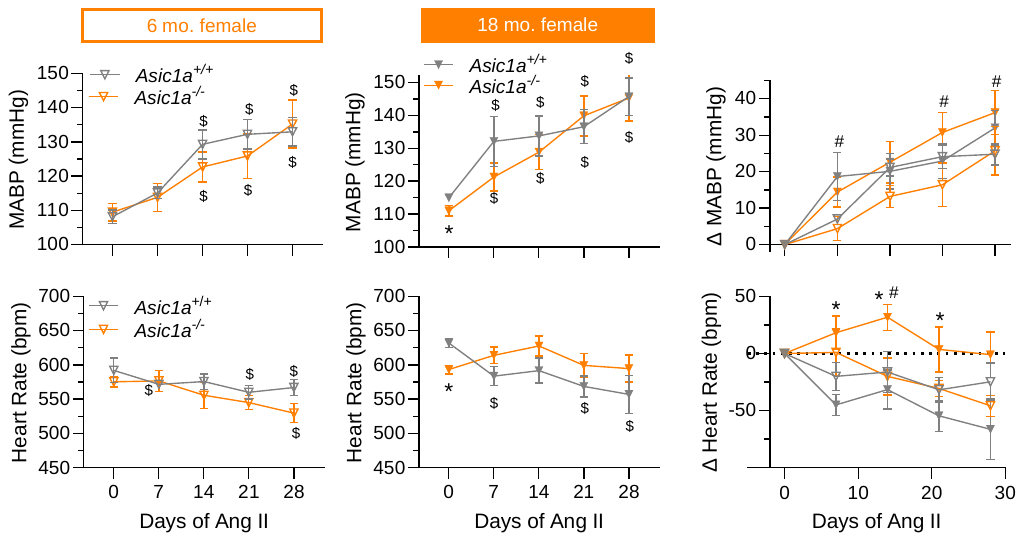
**Figure S2.** **Asic1a deletion leads to age-dependent protection against Ang II-induced hypertension in male mice. A,B**) MABP and **C**) changes in MABP in response to angiotensin II infusion (600ng/kg/min) over 28 days in A) adult (6-month-old) and B) aged (18-month-old) female mice. **D,E**) HR and **F**) changes in HR over the same time. n =5-6 animals per group; * p < 0.05 versus *Asic1a*^+/+^; $ p<0.05 vs time 0; # p<0.05 vs 6-month-old adult; analyzed by repeated measures two-way ANOVA, individual groups compared using Tukey’s multiple comparisons test.

**Figure S3.** Urine Epinephrine (Epi; ng/ml):Creatinine (mg/dl) ratio in 6- and 18-month-old *Asic1a*^+/+^ and *Asic1a*^-/-^ **A**) male and **B**) female mice. There were no significant differences in Creatinine (mg/dl) levels (**C,D**). Analyzed by two-way ANOVA.

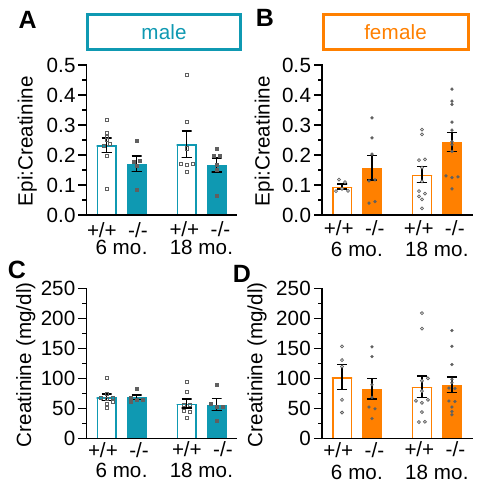

**Figure S4.** **Chemoreceptor sensitivity is intact in aged *Asic1a*^-/-^ mice.** Whole-body plethysmography was used to determine **A**) respiratory frequency (breaths/min) and **B**) tidal volume (μL/breath/g) from which **C**) minute ventilation (ml/min/g body wt) was calculated. Measurements were taken at baseline and following acute exposure to hypercapnia (6% CO_2_) or isocapnic hypoxia (7.0% O_2_, 3.2% CO_2_, balance N_2_) in 18-month-old male *Asic1a*^+/+^ and *Asic1a*^-/-^ mice. Values are means ± SEM; n = number of animals per group, analyzed by repeated measures two-way ANOVA, individual groups compared using Šídák's post-hoc test.

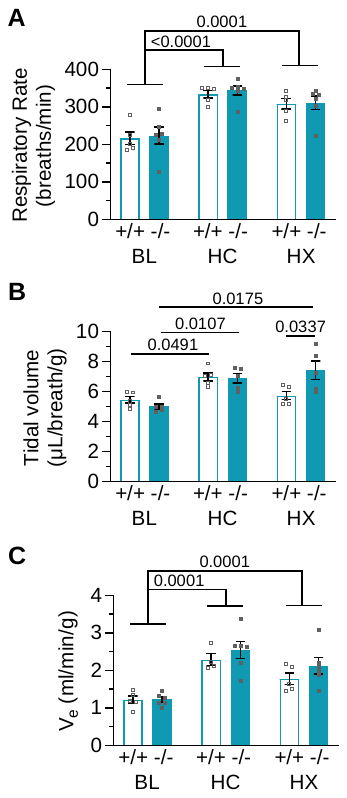

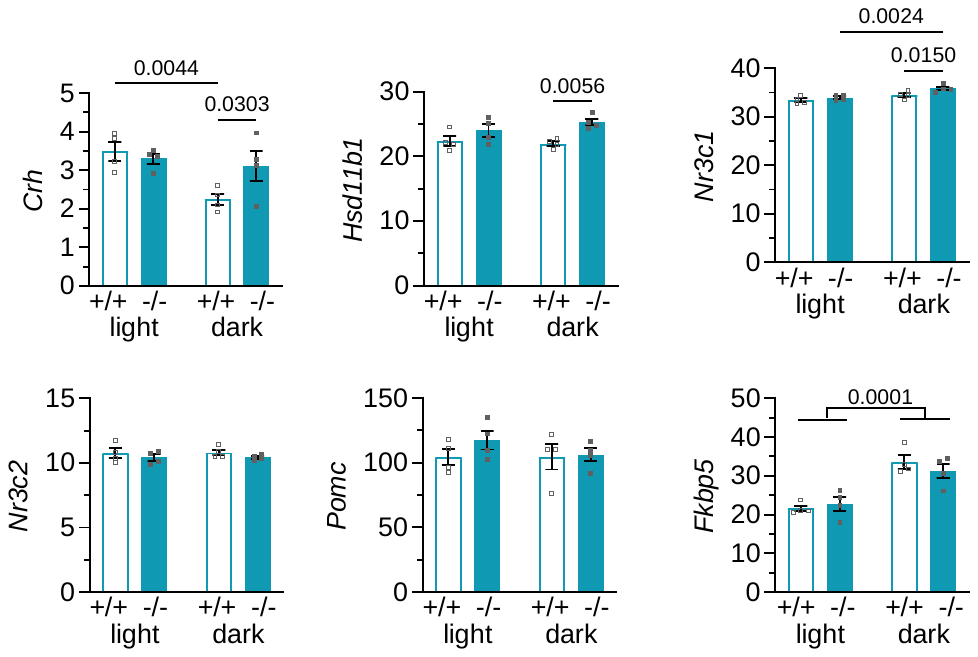
**Figure S5.** **ASIC1a Deficiency Alters Hypothalamic Gene Expression and Activates chronic stress pathway.** Transcripts per million (TPM) values (mean ± SEM) for individual genes retrieved from the Gene Expression Omnibus dataset GSE185178, which is an RNA-Seq dataset of hypothalami collected during the light and dark cycle from 3-month-old *Asic1a^+/+^* and *Asic1a^-/-^* (n=4) mice. Analyzed by two-way ANOVA, individual groups were compared using Fisher’s LSD. Corticotropin-releasing hormone (*Crh*); 11β-Hydroxysteroid dehydrogenase type 1 (*Hsd11b1*); glucocorticoid receptor (*Nr3r1*); mineralocorticoid receptor (*Nr3c2*); Proopiomelanocortin (*Pomc*); FK506-Binding Protein 5 (*Fkbp5*)

**Figure S6.** **Aged male *Asic1a*^-/-^ mice exhibit increased cardiac and coronary artery fibrosis. A**) cardiac fibrosis (% of 3.5 mm^2^ area). **B**) AZAN trichrome-stained heart sections and coronary arterial fibrosis in 6- and 18-month-old, male and female, *Asic1a*^+/+^ and *Asic1a*^-/-^ mice. AZAN trichrome shows cell nuclei (dark red), collagen (blue), and orange-red in cytoplasm. Values are mean ± SEM; n= animals/group; analyzed by two-way ANOVA, individual groups compared using Tukey’s multiple comparisons test.

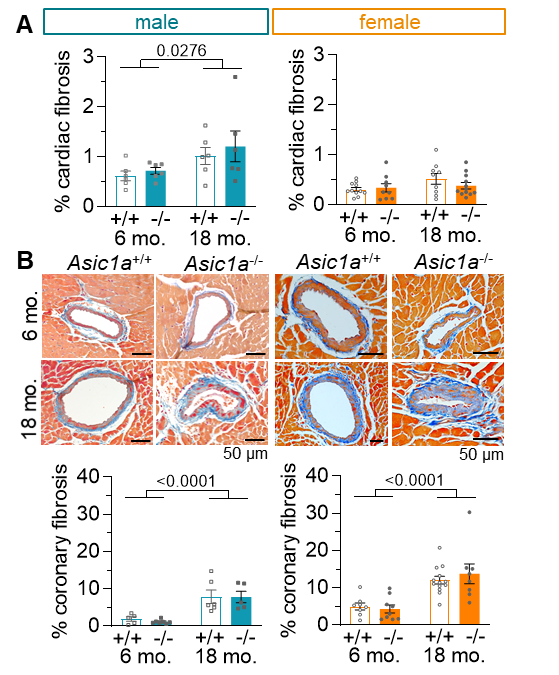

**Figure S7.** **In female mice, *Asic1a* deletion does not significantly alter kidney structure. A**) Representative AZAN trichrome-stained kidney sections and summary data showing B) percent renal fibrosis (% of 3.5 mm^2^ area) **C**) glomerular cross-sectional area (CSA; µm^2^ x 1,000), and **D**) glomeruli number (3.5 mm^2^ area). Measurements for urine **E**) osmolality (Osm/kg H2O) and **F**) protein (µg/µL) from 6- and 18-month-old *Asic1a*^+/+^ and *Asic1a*^-/-^ female mice. Values are mean ± SEM; analyzed by two-way ANOVA, individual groups compared using Tukey’s multiple comparisons test.

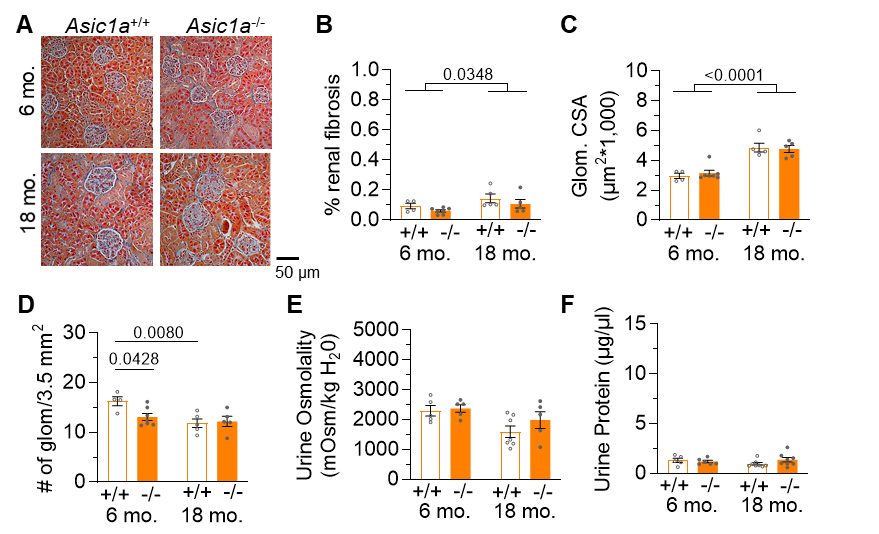
